# Metabolic and haemodynamic resting-state connectivity of the human brain: a high-temporal resolution simultaneous BOLD-fMRI and FDG-fPET multimodality study

**DOI:** 10.1101/2020.05.01.071662

**Authors:** Sharna D Jamadar, Phillip GD Ward, Emma Xingwen Liang, Edwina R Orchard, Zhaolin Chen, Gary F Egan

## Abstract

Simultaneous FDG-PET/fMRI ([18F]-fluorodeoxyglucose positron emission tomography functional magnetic resonance imaging) provides the capacity to image two sources of energetic dynamics in the brain – glucose metabolism and haemodynamic response. Functional fMRI connectivity has been enormously useful for characterising interactions between distributed brain networks in humans. Metabolic connectivity based on static FDG-PET has been proposed as a biomarker for neurological disease; but static FDG-PET cannot be used to estimate subject-level measures of *connectivity*, only across-subject *covariance*. Here, we applied high-temporal resolution constant infusion fPET to measure subject-level metabolic connectivity simultaneously with fMRI connectivity. fPET metabolic connectivity was characterised by fronto-parietal connectivity within and between hemispheres. fPET metabolic connectivity showed moderate similarity with fMRI primarily in superior cortex and frontoparietal regions. Significantly, fPET metabolic connectivity showed little similarity with static FDG-PET metabolic covariance, indicating that metabolic brain connectivity is a non-ergodic process whereby individual brain connectivity cannot be inferred from group level metabolic covariance. Our results highlight the complementary strengths of fPET and fMRI in measuring the intrinsic connectivity of the brain, and open up the opportunity for novel fundamental studies of human brain connectivity as well as multi-modality biomarkers of neurological diseases.

---

The invention of positron emission tomography (PET) in the 1970’s (Ter-Pogossian, 1992) and the development of functional magnetic resonance imaging in the early 1990’s (Ogawa et al., 1990) have provided unique insights into brain function in living humans. During the past two decades functional magnetic resonance imaging (fMRI) has been an enormously powerful tool for the discovery of intrinsic brain networks, the characterisation of brain connectivity, and the identification of interactions between distributed brain networks (Fox et al., 2007; Fox and Raichle, 2007). Recently, the advent of simultaneous [18F]-fluorodeoxyglucose PET (FDG-PET) and fMRI of the brain (Judenhofer et al., 2008) has provided the opportunity for unique new insights into the mechanisms of dynamic metabolic and neurovascular activity in the human brain (Villien et al., 2014).

Resting-state functional MRI measures the temporal coherence of spontaneous neural activity between spatially distinct brain regions (Fox and Raichle, 2007), and is normally measured using blood oxygenation level dependent fMRI (BOLD-fMRI) (Biswal et al., 1995). Resting-state brain connectivity indexes spontaneous large-amplitude changes in blood oxygenation at low frequencies that have been associated with variability in cognition (Fox et al., 2007; Jamadar et al., 2016), individual differences in age (Andrews-Hanna et al., 2007), sex (Jamadar et al., 2018), and psychiatric and neurological conditions (8). Resting-state fMRI connectivity has potential as a biomarker of clinical progression for a number of neurological diseases, although its application as a diagnostic biomarker is currently limited (Hohenfeld et al., 2018).

BOLD-fMRI provides a haemodynamic-based surrogate index of neuronal activity with a temporal resolution in the order of seconds and sub-millimeter spatial resolution. The BOLD-fMRI signal is an indirect index of neuronal function, arising from neurovascular coupling between neuronal activity and cerebral haemodynamics (Phillips et al., 2016). While BOLD-fMRI is usually interpreted as arising from neuronal activity, there are a number of non-neuronal contributors to the BOLD signal, including heart rate variability, respiration, head movement, individual variability in haemoglobin and the oxygen-carrying capacity of the blood, etc. (Liu, 2017; Ward et al., 2019). As such, BOLD-fMRI is a semi-quantitative non-absolute index of neural activity, and BOLD-fMRI responses cannot be quantitatively compared across brain regions, subjects, or imaging sites, or within the same individual across time (Logothetis, 2008). These characteristics are a major limiting factor in the development of BOLD-fMRI-based biomarkers, particularly when derived from fMRI functional connectivity metrics (Hohenfeld et al., 2018).

Resting-state connectivity measured using other neuroimaging techniques provide a unique perspective on information transfer in the brain. Here, we investigate brain connectivity derived from FDG-PET to measure co-variation of glucose uptake throughout the human brain; an index of cerebral metabolism. FDG-PET provides a snapshot of cerebral glucose uptake, and connectivity analyses of FDG-PET have characterised the covariation in cerebral glucose uptake across subjects (Horwitz et al., 1984; Moeller et al., 1987). This measure of ‘metabolic’ connectivity is an important complement to BOLD-fMRI functional connectivity, since FDG-PET represents a more direct and quantitative measure of neuronal function than BOLD-fMRI. While evidence suggests metabolic connectivity shows moderate-to-strong correlation with fMRI connectivity (Di et al., 2017; Di and Biswal, 2012; Passow et al., 2015; Savio et al., 2017), due to technical limitations, the majority of FDG-PET studies have acquired static images of the brain with an effective temporal resolution of the scan duration –10-40mins. Since static FDG-PET (sPET) acquisitions provide a single cumulative measurement per subject, metabolic ‘connectivity’ was estimated *across-subjects*. As such, these results are more accurately characterised as metabolic *covariance*, rather than *connectivity*. Covariance measures are not dependent on the temporal correlation of glucose uptake and therefore provide patterns that do not necessarily arise from coupled activity between brain regions. Furthermore, it is known from fMRI that group-level covariance poorly predicts individual measures of seed-based functional connectivity (Roberts et al., 2016). Consequently, static FDG-PET images cannot be used to estimate metabolic connectivity within single subjects, greatly limiting their application as a disease biomarker (Veronese et al., 2019; c.f., Yakushev et al., 2017).

Recent advances in radiotracer delivery, together with the improved PET signal detection sensitivity of dual-modality MR-PET scanners, has made it possible to study the dynamics of FDG-PET glucose uptake with substantially improved temporal resolution. The method described as ‘functional’ FDG-PET (FDG-fPET) have achieved a temporal resolution of 60sec (Hahn et al., 2016, 2017; Jamadar et al., 2019b; Li et al., 2019; Villien et al., 2014) or less (Hahn et al., 2020; Jamadar et al., 2019a; Rischka et al., 2018). These methodological advancements have opened up an unprecedented opportunity to examine the temporal coherence of glucose metabolic signals across subjects, in a similar manner to BOLD-fMRI.

The goal of the present study was to investigate FDG-fPET metabolic connectivity with high temporal resolution. We used FDG-fPET data with temporal resolution of 16sec, and simultaneously acquired BOLD-fMRI data with a temporal resolution of 2.45sec. Two major hypotheses were tested in the present study. Firstly, we hypothesised that static FDG-PET metabolic covariance would be moderately associated with fPET metabolic brain connectivity. Secondly, we predicted that resting-state metabolic brain connectivity would be strongly associated with restingstate BOLD-fMRI connectivity.

## Materials and Methods

All methods were reviewed by the Monash University Human Research Ethics Committee, in accordance with the Australian National Statement on Ethical Conduct in Human Research (2007). Administration of ionising radiation was approved by the Monash Health Principal Medical Physicist, in accordance with the Australian Radiation Protection and Nuclear Safety Agency Code of Practice (2005). For participants aged over 18yrs, the annual radiation exposure limit of 5mSv applies and the effective dose equivalent was 4.9mSv. Detailed information on the constant infusion acquisition procedure is available in Jamadar et al. (2019b). Data is available on OpenNeuro (see Data Availability Statement), and a data descriptor is available in Jamadar et al. (in press).

### Participants

Participants (n=27) were aged 18-23 years (mean 19 years); 20 female, all right handed (Edinburgh Handedness Inventory). Participants had between 13-18 years of education (mean 14 years), normal or corrected-to-normal vision and no personal history of diagnosed Axis-1 mental illness, diabetes or cardiovascular illness. Participants were screened for claustrophobia, non-MR compatible implants, clinical or research PET scan in the past 12-months, and women were screened for current or suspected pregnancy. Prior to the scan, participants were directed to consume a high protein/low sugar diet for 24-hours, fast for 6 hours, and drink 2-6 glasses of water. Blood sugar level was measured using an Accu-Check Performa (model NC, Mannheim, Germany); all participants had blood sugar levels below 10mmol/L (max in this sample 4.73mmol/L).

### Procedure

Prior to the scan, participants completed a brief cognitive battery (30mins, results not reported here). Participants were then cannulated in the vein in each forearm with a minimum size 22-gauge cannula, and a 10mL baseline blood sample was taken at the time of cannulation. For all participants, the left cannula was used for FDG infusion, and the right cannula was used for blood sampling. Primed extension tubing was connected to the right cannula (for blood sampling) via a three-way tap.

Participants underwent a 95-minute simultaneous MR-PET scan in a Siemens (Erlangen) Biograph 3 Tesla molecular MR (mMR) scanner (Figure 1A). Participants were positioned supine in the scanner bore with head in a 16-channel radiofrequency (RF) head coil, and were instructed to lie as still as possible with eyes open, and think of nothing in particular. [18-F] fluroodeoxyglucose (FDG; average dose 233MBq) was infused over the course of the scan at a rate of 36mL/hr using a BodyGuard 323 MR-compatible infusion pump (Caesarea Medical Electronics, Caesarea, Israel). One participant received a lower dose (167MBq) due to infusion pump error. Infusion onset was locked to the onset of the PET scan.

**Figure 1:**
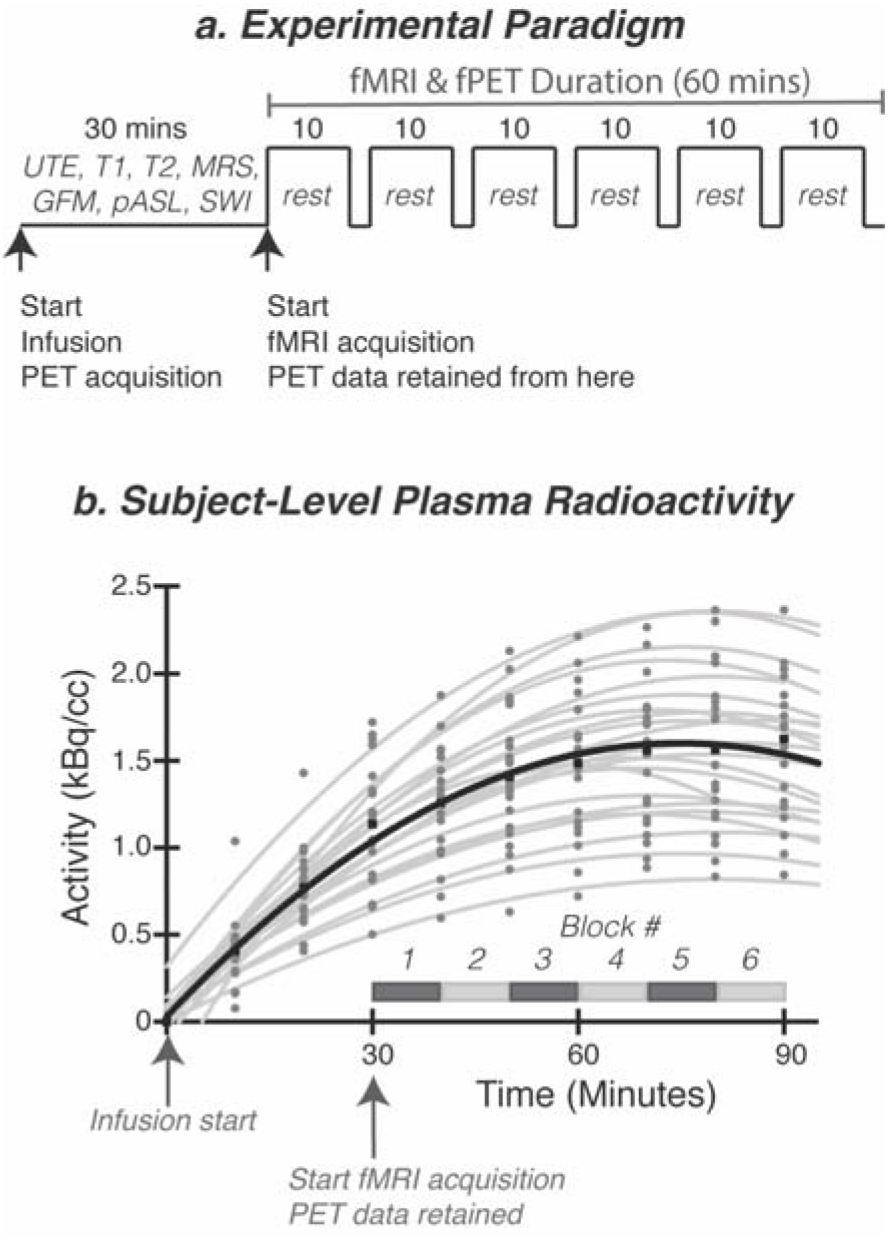
**A**. Experimental paradigm. Participants underwent a 95-minute simultaneous MRI-PET scan. [18F]-FDG was infused over the course of the scan, with infusion start time locked to the onset of the PET scan (time 0). For the first 30-minutes, non-functional MRI scans were acquired to allow the PET signal to increase to detectable levels (see Figure 2, plasma radioactivity curve). At the 30-minute timepoint, 6 consecutive 10-minute resting-state BOLD-fMRI (T2* echo planar imaging) blocks were completed. Participants rested quietly with eyes open with central fixation (cross) for the full 60-minutes. Abbreviations: UTE, ultrashort echo time; MRS, magnetic resonance spectroscopy; GFM, gradient field map; pASL, pulsed arterial spin labelling; SWI, susceptibility weighted imaging; mins, minutes. **B.** Plasma radioactivity curve. Individual subject radioactivity curves are plotted in grey, with the average across all subjects plotted in black.

Plasma radioactivity levels were measured throughout the duration of the scan. At 10-mins post-infusion onset, a 10mL blood sample was taken from the right forearm using a vacutainer; the time of the 5mL mark was noted for subsequent decay correction. Subsequent blood samples were taken at 10-min intervals for a total of 10 samples for the duration of the scan. The cannula line was flushed with 10mL of saline after every sample to minimise line clotting. Immediately following blood sampling, the sample was placed in a Heraeus Megafuge 16 centrifuge (ThermoFisher Scientific, Osterode, Germany) and spun at 2000rpm for 5 mins; 1000 μL plasma was pipetted, transferred to a counting tube and placed in a well counter for 4mins. The count start time, total number of counts, and counts per minute were recorded for each sample. Figure 1B shows the interpolated plasma radioactivity concentration over time. The average radioactivity concentration constantly increases over time with the lowest relative slope at the end of the acquisition.

### MR-PET protocol

PET data was acquired in list mode. Infusion of FDG radiotracer and PET data acquisition started with the Ultrashort TE (UTE) MRI for PET attenuation correction (Figure 1A). To allow the PET signal to rise to detectable levels, non-functional MRI scans were acquired in the first 30-mins following infusion onset. These scans, in order, were: UTE (TA=1.40mins), T1 3D MPRAGE (TA = 7.01mins, TR=1640ms, TE=2.34ms, flip angle = 8°, FOV=256×256mm^2^, voxel size = 1×1×1mm^3^, 176 slices; sagittal acquisition), and several scans not reported here: T2 SPACE (TA=5.52min), magnetic resonance spectroscopy (MRS, TA=2.48min), gradient field map (TA=1.02min), pulsed arterial spin labelling (TA=4.21), T2 susceptibility-weighted image (TA=6.50min), left-right phase correction (TA=0.21min). For the remainder of the scan, six consecutive 10min blocks of T2*-weighted echo planar images (EPIs) were acquired (TR=2450ms, TE=30ms, FOV=190mm, 3×3×3mm3 voxels, 44 slices, ascending axial acquisition).

### MR image preparation

For the T1 structural image, the brain was extracted, then registered to MNI152 space using Advanced Normalization Tools (ANTs) (Avants et al., 2011). The grey matter, white matter and brain cortex labels of T1 image were segmented by using Freesurfer with Desikan-Killiany Atlas (Diedrichsen et al. 2009).

The six blocks of EPI scans for all subjects (a total of 245 EPIs) underwent a standard fMRI pre-processing pipeline. Specifically, all scans were brain extracted (FSL BET (Smith, 2002)), corrected for intensity non-uniformity using N4 bias field correction (ANTs (Tustison et al., 2010)), motion corrected (FSL MCFLIRT (Jenkinson et al., 2002)), slice timing corrected (AFNI (Cox, 1996) and high-pass filtered (Hz > 0.01) to remove low frequency noise (FSL (Jenkinson et al., 2012). Across subjects, average mean framewise translational motion was 0.41mm; maximum was 1.09mm.

### PET image reconstruction and preparation

The 5700-second list-mode PET data for each subject was binned into 356 3D sinogram frames each of 16-second interval. The pseudoCT method (Burgos et al., 2014) was used to correct the attenuation for all acquired data. Ordinary Poisson-Ordered Subset Expectation Maximization (OP-OSEM) algorithm (3 iterations, 21 subsets) with point spread function correction was used to reconstruct 3D volumes from the sinogram frames. The reconstructed DICOM slices were converted to NIFTI format with size 344×344×127 (voxel size: 2.09 × 2.09 ×2.03 mm^3^) for each volume. A 5-mm FWHM Gaussian post-filter was applied to each 3D volume. All 3D volumes were temporally concatenated to form a 4D (344 × 344 × 127 × 356) NIFTI volume. A guided motion correction method using simultaneously acquired MRI images was applied to correct the motion during the PET scan. We retained the 225 16-sec volumes commencing from the 30 minute timepoint, which matched the start of the BOLD-fMRI EPI acquisition, for further analyses. A single static PET image was derived from the sum of the 16-sec volumes.

The 225 PET volumes were motion corrected (FSL MCFLIRT (Jenkinson et al., 2002); the mean PET image was brain extracted and used to mask the 4D data. The fPET data was further processed using a spatio-temporal gradient filter to estimate the short-term change in glucose uptake from the cumulative glucose uptake that was measured. The filter removed the accumulating effect of the radio-tracer and other low-frequency components of the signal to isolate short-term resting-state fluctuations. This approach intrinsically adjusted for the mean signal whilst avoiding global-signal regression and other approaches that may create spurious anticorrelations in the data (Li et al., 2019); (Murphy and Fox, 2017). Due to radiotracer dynamics, it was not expected that the fPET sensitivity would be uniform across the 60minutes of the resting-state data acquisition. As the radiotracer accumulated in the brain, it was anticipated that the signal-to-noise ratio of the PET image reconstruction would progressively improve. The Supplement includes the definition of the spatiotemporal filter (Supplementary Figure S1).

### Connectivity Analyses

fPET and fMRI timeseries were extracted for each of the 82 regions of interest from the segmentation of the T1-weighted image, interpolated using an ANTs rigid registration (Avants et al., 2011). To construct a connectivity matrix, Pearson’s correlation coefficients were estimated between the timeseries from pairs of regions. This produced a single per-subject per-modality 82×82 matrix corresponding to the 60minutes of resting-state in the experimental protocol. The six motion parameters were used to account for framewise displacement effects in the fPET and fMRI connectivity matrices. Thus, each region-by-region association is a partial correlation of region A, region B, pitch, roll, yaw, x, y, z for each subject, then averaged across subjects for the group connectome.

#### Group-average connectivity for BOLD-fMRI and FDG-fPET

The similarity between the group-average BOLD-fMRI and FDG-fPET matrices was assessed using a Pearson correlation across all edges on the lower triangle of the connectivity matrices (excluding the diagonal and symmetric triangle). The regional variation between the two modalities was assessed by calculating a separate correlation coefficient for each row of the connectivity matrix, providing a measure of similarity of the connectivity profile for each brain region.

#### sPET metabolic covariance

A FDG-sPET metabolic covariance matrix was constructed using the mean signal for each subject from the static PET image. The signal was de-meaned for each subject to remove inter-subject variances, such as dosage and physiology. The across-subject correlation between pairs of ROIs was then estimated to generate the covariance matrix. The similarity of across-subject metabolic covariance of sPET and inter-subject temporal correlation/metabolic connectivity of fPET was examined at a regional and global level. A comparison of the characteristics of the metabolic covariance and the metabolic connectivity was performed, assessing the scale of correlations and the network profile.

#### Spatial maps of region degree

To further characterise and contrast connectivity from the two modalities, the strength of connections to each region were examined. Each connectivity matrix was discretised at the 90^th^ percentile level to provide a binary graph of the ‘strongest’ connections. We took this approach, rather than a threshold approach based on null-hypothesis testing, as the SNR properties of each modality are substantially different. Thus, a common threshold across all three modalities would lead to many significant regions in some modalities (i.e. sPET) and very few in others (i.e. fPET). We argue that the percentile approach is appropriate in this analysis, as the goal was to examine and compare the distribution of most connected regions between modalities, without the additional concerns of subthreshold results. Results across other thresholds are reported in the Supplement. The number of these binary edges connected to each region was calculated to provide a map of regional degree for each modality.

#### Graph-based network analysis

To investigate the patterns of connectivity within and between subnetworks, node degree was calculated and regions were sorted by subnetwork assignment (i.e., frontoparietal, dorsal attention, ventral attention, default mode, somatomotor, limbic, subcortical, and visual; as classified by (Yeo et al., 2011). As the regions were not evenly distributed between subnetworks, some subnetworks (e.g. default mode network) contained a higher number of nodes than others. Therefore, there was a higher chance that significant edges would occur within subnetworks with more nodes by random chance. To minimise this bias in our interpretation, we adjusted node degree for the capacity of each network (number of observed edges, divided by number of potential edges between two subnetworks).

#### Shared variance between sPET & fPET; and fMRI & fPET

To investigate the shared variance between the two modalities, the fPET connectome for each individual was regressed from the sPET and fMRI connectome, respectively. The standardised residuals of the regression were visualised in a matrix to show the distribution of variance not explained by fPET in the sPET and fMRI connectome.

## Results

We first provide a qualitative overview of the static PET (sPET) covariance, functional PET (fPET) connectivity, and fMRI connectivity matrices (Figure 2), and their associated network graphs (Figure 3). Secondly, we compare the sPET results to fPET, to determine if sPET covariance is predictive of fPET connectivity. Lastly, we compare fPET results to fMRI, to determine the relationship between metabolic and functional connectivity.

**Figure 2:**
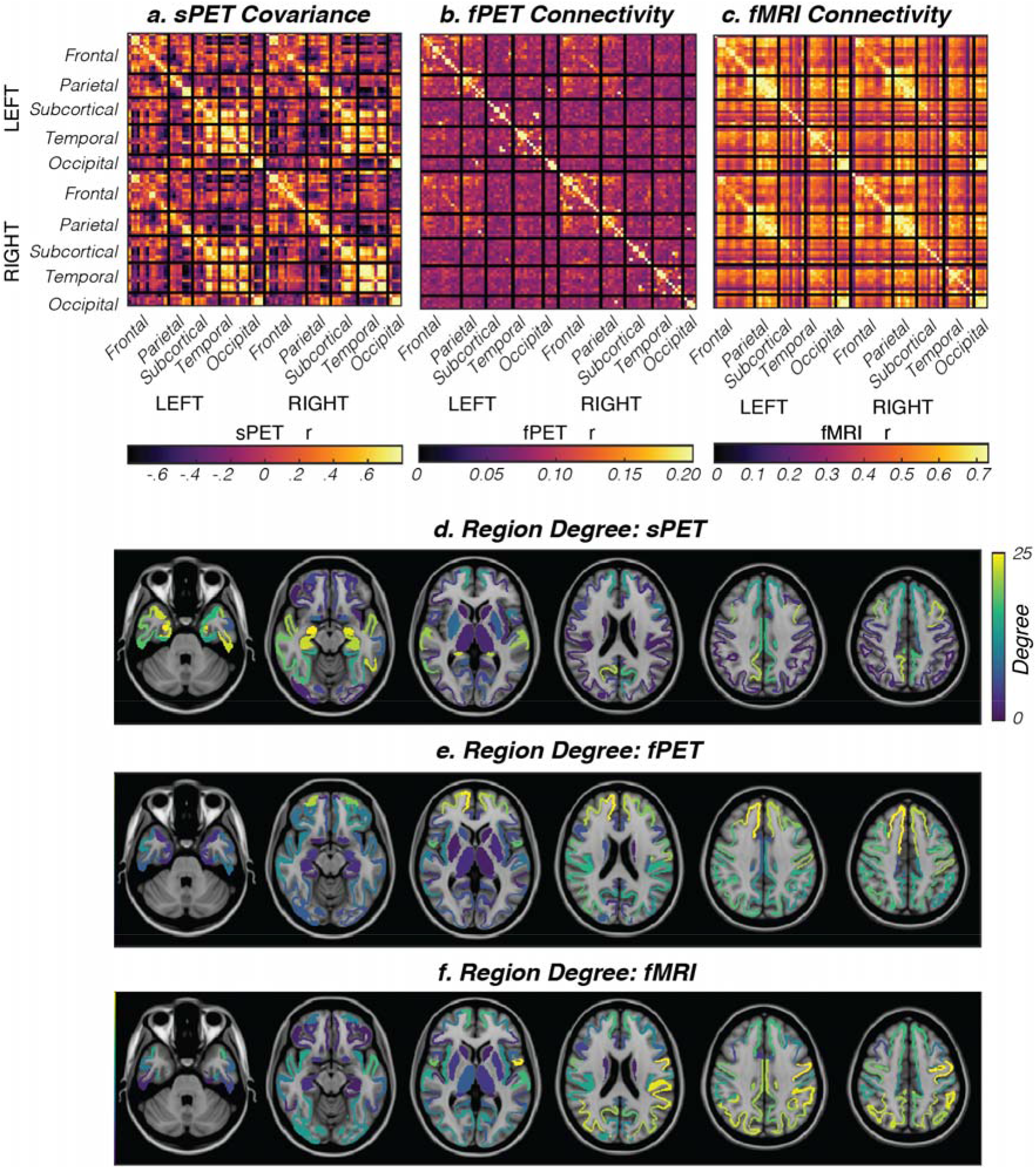
**A**. FDG-sPET resting-state covariance matrix thresholded from −0.65 < r < 0.65. **B.** FDG-fPET resting-state connectivity matrix thresholded from 0 < r < 0.12. **C.** BOLD-fMRI resting-state connectivity matrix thresholded from 0 < r < 0.75. Supplementary Figure 1 shows the distribution of the number of edges across r values for each modality. **D**. Spatial representation of the connection centrality (region degree; number of connections attached to the region) of FDG-sPET. **E.** Spatial representation of the connection centrality (region degree) of FDG-fPET and **F.** Spatial representation of the connection centrality (region degree) of BOLD-fMRI. Supplementary Figures 5–7 show the variation of the region degree plots (panels d-f) across different thresholds.

**Figure 3:**
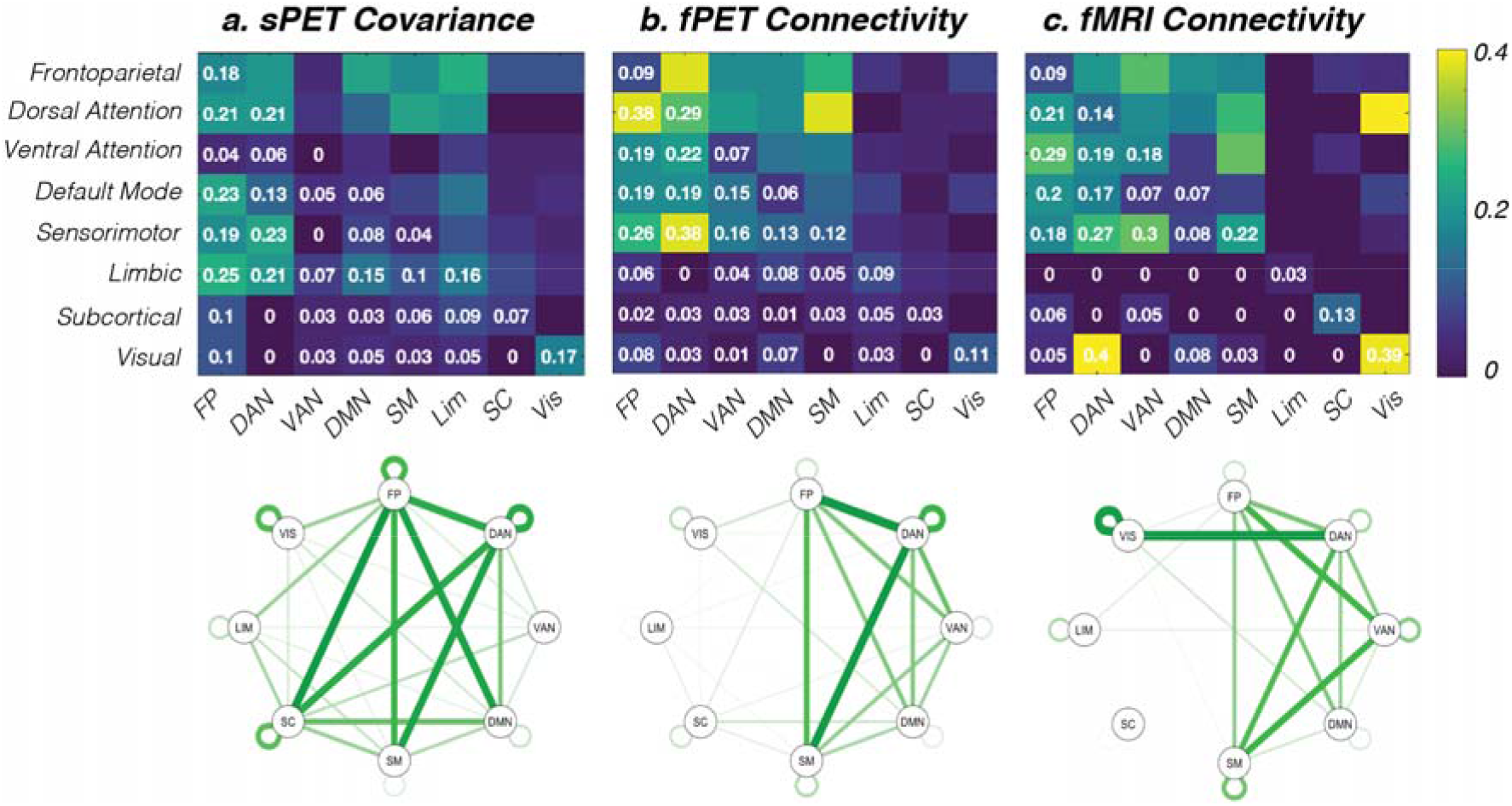
Connectivity matrices (top panel) and network graphs (bottom panel) for eight canonical resting-state networks. Within-network connections lie on the diagonal (e.g. visual-visual), and between-network connections lie on the off-diagonal (e.g. visual-default mode). **A.** FDG-sPET metabolic covariance, **B.** FDG-fPET metabolic connectivity and **C.** BOLD-fMRI haemodynamic connectivity. Abbreviations: FP, frontoparietal network; DAN, dorsal attention network; VAN, ventral attention network; DMN, default mode network; SM, sensorimotor network; Lim, Limbic network; SC, subcortical network; Vis, visual network.

### Metabolic covariance, metabolic connectivity, and haemodynamic connectivity

#### sPET Metabolic Covariance

The across-subject sPET metabolic covariance matrix showed large variability in connection strength across ROIs (Figure 2A). Each cortical sub-division showed strong covariance with neighbouring regions, as evident by the high covariance values (r≥0.5; orange-yellow colours) along the main diagonal. Furthermore, each cortical sub-division showed strong homotopic covariance, as evident by the high covariance values along the diagonals of the top right and bottom left quadrants of the matrix. High temporo-subcortical covariance was observed both within and between hemispheres; and high fronto-subcortical, fronto-temporal and parieto-occipital covariance was also apparent albeit of smaller magnitude than the temporo-subcortical covariance. FDG-sPET showed the highest region degree in the temporal poles and subcortically (Figure 2D). In the network graphs (Figure 3A), sPET showed strong widespread connectivity between the dorsal attention, frontoparietal, subcortical, sensorimotor and default mode networks. *fPET Metabolic Connectivity:* The most salient effect for the group-averaged within-subject FDG-fPET metabolic connectivity matrix was the stronger (r≥0.15; orange-yellow colours) fronto-parietal connectivity, both within and between hemispheres (Figure 2B). Left-right homotopic connectivity was also evident for frontal and parietal cortices; this was not visually apparent for subcortical, temporal and occipital cortices. fPET showed the highest degree in the frontal poles and the superior cortex (Figure 2E). For the network graphs (Figure 3B) fPET showed high connectivity between the frontoparietal and dorsal attention networks, and between the dorsal attention and sensorimotor networks. Moderate connectivity was apparent between the sensorimotor-frontoparietal networks, and between the default mode and dorsal attention, frontoparietal and visual attention networks. The dorsal attention network showed strong within-network connectivity as well.

#### fMRI Functional Connectivity

BOLD-fMRI showed strong (r≥0.6; orange-yellow colours) connectivity within anatomical sub-divisions, both within and between hemispheres (Figure 2C). This pattern was evident from the four diagonal lines in the fMRI connectivity matrix. A number of long-range connections between anatomical sub-divisions were also evident, including fronto-parietal, parieto-occipital and temporo-parietal regional connectivity. These long-range connections were evident both within and between hemispheres but were of smaller magnitude than the shortrange and homotopic connections. Network graphs for the fMRI data (Figure 3C) showed strongest connectivity between the dorsal attention and visual networks, with the visual network having the highest self-connectedness. The visual attention network showed high connectedness with the frontoparietal and sensorimotor network, and the default mode network showed an intermediate level of connectedness with the frontoparietal and dorsal attention networks. The anatomical projections of the most highly connected regions (Figure 2F, lower row) indicated that the parietal cortex was most inter-connected region in the fMRI data, with superior frontal and inferior occipital cortices showing intermediate levels of haemodynamic based connectivity. Subcortical and orbitofrontal regions were the least inter-connected regions in the BOLD-fMRI data.

### Comparison of FDG-sPET metabolic covariance and FDG-fPET metabolic connectivity

Qualitatively the sPET covariance and fPET connectivity matrices show little similarity (Figure 2A & 2B). The fPET connectivity was dominated by fronto-parietal connectivity, whereas the sPET covariance demonstrated greater variability throughout the connectivity matrix. The distribution of edges for each r value for sPET and fPET showed that fPET is biased towards positive r values, whereas sPET is symmetric around r=0 (Supplementary Figure S2). Comparison of the network graphs (Figure 3) and the anatomical projections of the most connected regions (Figure 2D-F) are consistent with the finding that sPET and fPET show little anatomical similarity across the brain. This result is confirmed by the matrix of sPET~fPET residuals (Figure 4A) which highlights that sPET covariance has substantial variance (e > 0.5) that is unexplained by fPET connectivity. In particular, the left-right homologue covariance in the subcortical and temporal regions is not explained by fPET within-subject connectivity. Taken together, these results are consistent with the conclusion that across-subject sPET connectivity does not predict within-subject fPET connectivity. These results are not consistent with the hypothesis that static metabolic connectivity would be moderately associated with within-subject metabolic brain connectivity.

**Figure 4:**
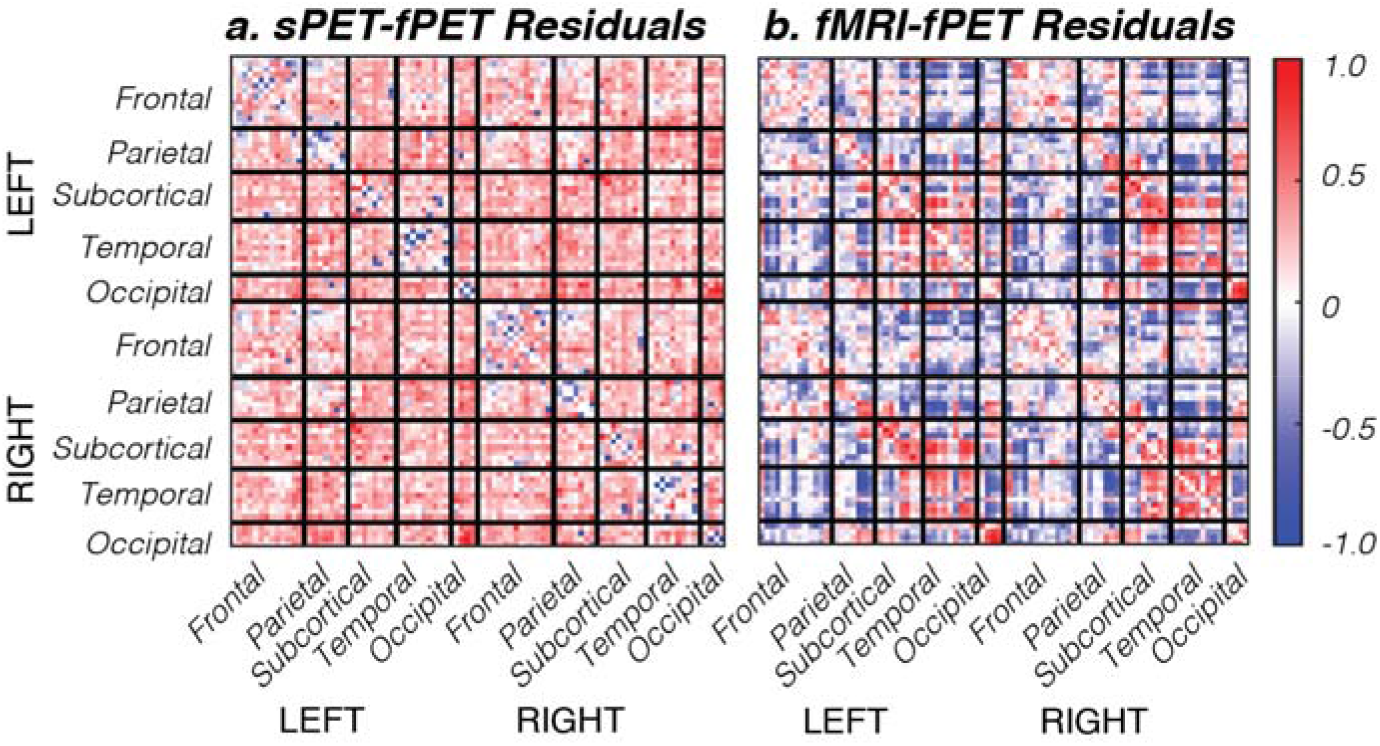
Matrix of residuals for the multiple regressions controlling for fPET variance for (**A**) sPET and (**B**) fMRI.

### Comparison of FDG-fPET metabolic connectivity and BOLD-fMRI haemodynamic connectivity

Qualitatively the FDG-fPET metabolic connectivity and BOLD-fMRI haemodynamic connectivity matrices showed moderate similarity, with a trend towards stronger connectivity associated with the fronto-parietal cortex (Figure 2). Both fPET and fMRI network graphs (Figure 3) showed strong connectivity for the dorsal attention network, with the dorsal attention network strongly connected to the frontoparietal network for fPET, and strongly connected to the visual network for fMRI. The default mode network showed moderate connectedness for both fPET and fMRI. The matrix of the fMRI~fPET residuals (Figure 4B) is relatively flat, with little variability across ROIs. Residual values were uniformly slightly positive (e ~0.2) suggesting a slight positive bias in variance for fMRI compared to fPET. Unlike the sPET~fPET residuals, there is little systematic variation across anatomical regions. These results are consistent with the conclusion that within-subject resting-state metabolic brain connectivity is more closely associated with resting-state haemodynamic connectivity than sPET metabolic covariance.

## Discussion

In this study, we report for the first time resting-state metabolic brain connectivity measured using a novel FDG-functional PET imaging protocol and methodology in humans. A key strength of this work is the simultaneous acquisition and high temporal resolution that enabled multi-modality within-subject analyses of resting state brain activity. The simultaneous acquisition of fPET and fMRI data is of particular importance, as measures of metabolic and haemodynamic responses to the same neural activity without the confound of intra-individual differences in attention, fatigue, motivation, nutrient intake and blood chemistry that occur in sequential brain imaging experiments (Chen et al., 2018). We compared shared variance between within-subject functional PET metabolic connectivity and across subject static PET metabolic covariance, and with BOLD-fMRI haemodynamic connectivity. Resting-state functional FDG-PET connectivity showed a high degree of unexplained variance between sPET covariance and fPET connectivity; and greater explained variance between fPET connectivity and BOLD-fMRI connectivity. Our study builds upon the foundations of earlier work that has used variants of positron emission tomography to study the resting brain, including the seminal work of Fox, Raichle, Shulman and colleagues who used O^15^ PET imaging of cerebral blood flow to delineate the default mode network (Raichle et al., 2001; Shulman et al., 1997), and Horwitz et al. (Azari et al., 1994; Horwitz et al., 1984) who conducted the first region-by-region correlation of neuroimaging signals using FDG-PET.

### Metabolic connectivity is predominantly evident in fronto-parietal regions

FDG-fPET metabolic connectivity was dominated by fronto-parietal connectivity both within and between hemispheres. This is consistent with early evidence showing the frontoparietal cortices are highly metabolically active compared to temporo-occipital regions (Sasaki et al., 1986), particularly at rest (Shokri-Kojori et al., 2019); (Vaishnavi et al., 2010). This pattern was confirmed by the brain network graphs where the dorsal attention network was strongly connected to the frontoparietal and sensorimotor networks, and showed strong intra-connectedness. Together, the frontoparietal and dorsal attention networks are responsible for flexible, goal directed behaviour (Laird et al., 2011), and the two networks interact closely to subserve action-focused attentional control (Dixon et al., 2018). The inter-connectedness of the frontoparietal and dorsal attention networks with the sensorimotor network is consistent with this interpretation. Interestingly, the dorsal attention and frontoparietal control networks have the highest metastability, update efficiency, and neural synchrony in comparison to other resting-state networks (Alderson et al., 2020). These networks demonstrate a resting-state configuration that is similar to a task-related configuration, which thereby facilitates dynamic and flexible switching between rest and task-active states. Our results support this interpretation and suggest that glucose metabolism in these networks is maintained at a high level in the resting-state, potentially to facilitate flexible switching to task-positive states when necessary.

The benefits of fPET methodology are twofold. Firstly, FDG uptake is neurophysiologically closer to the process of interest – i.e. neuronal function – than the BOLD response. It is well-known that BOLD-fMRI is an indirect index of neuronal activity that is not only ill-defined (e.g., (Phillips et al., 2016), and non-quantitative (Logothetis, 2008), but is also confounded by physiological parameters (e.g., (Ward et al., 2019). In contrast, most of the glucose uptake measured by FDG-PET is localised to the synapses (Sokoloff et al., 1977); (Sokoloff, 1981); (Magistretti and Allaman, 2015). As noted by (Raichle and Snyder, 2007), the use of PET methodologies in characterising the function of the brain at rest is important, as the quantitative nature and physiological underpinnings of the method allows one to uncouple effects of blood flow, oxygen consumption and glucose metabolism in the measured signal. Although technical considerations have led to the current situation where BOLD-fMRI is the more common method for visualising the spatiotemporal dynamics of neural function, FDG-PET is a more direct and less confounded method for visualising neuronal activity. The second benefit of fPET is that the method provides information about the dynamic use of the primary energy source of the brain — glucose. Univariate snapshots of glucose hypometabolism are predictive of neurodegeneration (e.g., (Jack et al., 2013), however preliminary evidence suggests that changes in the dynamic use of glucose over time may be an earlier and more powerful predictor of metabolic changes in neurodegenerative disease (Sanabria-Diaz et al., 2013). FDG-fPET methodology therefore represents an important development in characterising metabolic connectivity in the human brain and may show promise as a biomarker for disease in the future.

### Metabolic covariance is a poor predictor of metabolic connectivity

FDG-sPET metabolic covariance was dominated by left-right homologue, frontoparietal, temporo-subcortical and occipito-temporal covariance. This pattern is consistent with the earliest evidence of region-to-region correlation of the cerebral metabolic rate of glucose consumption (CMRGLC) reported by Horwitz and colleagues over 30-yrs ago (e.g., (Horwitz et al., 1984); (Azari et al., 1994). In their early work, Horwitz et al. noted the paucity of positive correlations between the occipito-temporal and fronto-parietal lobes; a finding which is evident in the present results where fronto-temporal and parieto-occipital edges demonstrated negative correlations. Our sPET metabolic covariance results are also consistent with more recent published findings. Using a static PET acquisition, (Di and Biswal, 2012) reported that independent component analysis (ICA) decomposition revealed resting-state networks predominantly between left-right homotopic regions, with few networks showing anterior-posterior connectivity. In their follow-up report, (Di et al., 2017) again found that sPET covariance is dominated by left-right homotopic connections, and short-range connections within anatomical sub-divisions, with only a few fronto-parietal connections. Similarly, (Savio et al., 2017) reported that sPET resting-state networks showed modest overlap with BOLD-fMRI networks, but were sparser and less well-separated from each other compared to BOLD networks (see also (Passow et al., 2015)). In summary, the static FDG-PET metabolic covariance results in the present study are consistent with the extensive literature of previous resting-state studies.

We compared fPET connectivity to sPET covariance, since across-subject sPET covariance has (until the present study) been the established method for examining metabolic ‘connectivity’ in health and disease (Yakushev et al., 2017). On the basis of previous comparisons of across-versus within-subject haemodynamic connectivity (Roberts et al., 2016), we hypothesised that sPET covariance would be moderately associated with fPET connectivity. Strikingly, we found that the sPET covariance matrix shows a high degree of unexplained variance when fPET connectivity was added to the model as a predictor. In other words, across-subject sPET covariance is a poor predictor of within-subject fPET connectivity. It is striking that the highest levels of unexplained variance was obtained in temporal and subcortical intra-hemisphere (left-left & right-right) and inter-hemisphere (left-right, right-left) connections. This suggests that the left-right homologue covariance identified previously (e.g., Di & Biswal, 2012; Di et al., 2017) may be an artifact of the across-subject correlation, and poorly reflective of within-subject connectivity. This somewhat surprising finding is analogous to Simpson’s Paradox (Simpson, 1951) whereby a statistical relationship observed at a population level is reversed at the level of the individuals that constitute the population (Kievit et al., 2013). Conceptually, Simpson’s Paradox refers to a sign reversal in the statistical relationships obtained at the individual and population (or group) levels. Roberts and co-workers found evidence of Simpson’s Paradox for a number of brain regions in fMRI haemodynamic connectivity in the dorsal attention and default mode networks (Roberts et al., 2016). However, they found that the majority of brain regions did not show a full reversal of the correlation values, as most brain regions showed across-subject correlations in the same direction as the within-subject correlations. Roberts et al. therefore concluded that across-subject fMRI covariance is a reasonable predictor of within-subject fMRI connectivity.

While we did not directly test for Simpson’s Paradox in our data, we found that there is a substantial amount of variability in the group-level sPET covariance matrix that was not explained by within-subject fPET connectivity. Simpson’s Paradox is a special case of the *ecological fallacy* (Robinson, 1950), and the related concept of *ergodicity* (Molenaar, 2008); (Kievit et al., 2013). Ergodic processes occur when a group-level result is generalisable to the individuals within the sample. In reality, ergodic processes are quite rare, because two strict criteria must be met: namely, the process must be homogeneous across individuals within a sample, and the statistical parameters that describe the process must be constant or stationary over time. Functional brain connectivity measured using neuroimaging data does not meet the criteria for ergodicity and thus group-level results are not generalisable to the individuals within the group (Fisher et al., 2018). Liegeois and colleagues have provided a complete mathematical examination of ergodicity in the context of BOLD-fMRI functional connectivity data ((Liégeois et al., 2017). Due to technical limitations of the PET imaging method, existing studies of resting-state metabolic connectivity has been limited to examinations of group-level covariance rather than within-subject correlation of FDG-PET timeseries (e.g., (Horwitz et al., 1984); (Di et al., 2017); (Savio et al., 2017). Researchers have attempted to draw quite strong conclusions on the basis of these results including attempts to use metabolic covariance as a biomarker for disease (Yakushev et al., 2017). A biomarker must, by definition, be estimable at the individual level (FDA-NIH, 2016). Our results suggest that metabolic covariance cannot be used to predict within-subject connectivity. While sPET covariance analyses may be useful in other contexts, future attempts to explore metabolic connectivity as a biomarker for disease must necessarily use fPET as the only statistically valid approach.

### FDG-fPET metabolic connectivity is correlated with BOLD-fMRI haemodynamic connectivity in fronto-parietal regions

After controlling for FDG-fPET metabolic connectivity, the BOLD-fMRI connectivity matrix was uniformly slightly positive, with little variation across regions. In other words, fPET and fMRI share a greater amount of variance by comparison to fPET and sPET. The similarity between fPET and fMRI was also apparent when qualitatively comparing the network graphs and the distribution of the regions with higher degrees of connectivity. These results complement the recent findings of (Amend et al., 2019) who reported that FDG-fPET metabolic connectivity showed similarities to BOLD-fMRI haemodynamic connectivity in a rodent model. Our findings are also consistent with recent results of (Shokri-Kojori et al., 2019) who found that frontoparietal regions had the highest concurrent energy use (based on CMR_GLC_) and BOLD-fMRI activity, in comparison to the rest of the brain.

One puzzling finding was the low degree apparent in subcortical regions for both FDG-fPET and BOLD-fMRI. Indeed, this effect was also apparent for the FDG-sPET results. These observations are in contrast with the known high degree of interconnectedness of the subcortical regions with the rest of the brain: the cortico-basal ganglia-cerebellar (Middleton and Strick, 2000); (Bostan and Strick, 2010) and cortico-thalamic circuits (Parent and Hazrati, 1995). These results are likely attributable to the reduced sensitivity of signal detection in midbrain areas for both PET and fMRI (Supplementary Figure S3). Smaller subcortical structures are also likely to be subject to partial volume effects (Hoffman et al., 1979), particularly in PET data which has a physical limit in spatial resolution due to the positron range (Moses, 2011), which may also contribute to this result. Reconstruction techniques to improve PET spatial resolution, and post-processing techniques to reduce partial volume effects (Sudarshan et al., 2020) may result in higher quality and greater brain coverage of the fPET connectivity matrices.

Direct comparisons between FDG-fPET and BOLD-fMRI are in part limited by the differences in temporal resolution and signal-to-noise ratio (SNR) of the two imaging modalities. The FDG-fPET protocol involves the administration of low levels of radioactivity over an extended time period, and results in PET images with lower signal and greater noise in comparison to BOLD-fMRI. Neural activity is optimally investigated using imaging techniques that have temporal resolutions in the order of milliseconds, which cannot be achieved by either fPET or fMRI. Nevertheless, the temporal resolution of conventional fMRI has been enormously useful for characterising distributed brain activity involving networks with multiple synapses over large-scale pathways in the brain. Importantly, although our understanding of integrative human brain networks has been developed largely from fMRI experimental results, the technique cannot serve as a ‘ground-truth’ for brain networks that are identifiable using FDG-fPET. In addition to the inherent methodological limitations of fMRI, the FDG-fPET and BOLD-fMRI experimental techniques detect different physiological targets; namely neural glucose metabolism and deoxygenation of haemoglobin in the cerebrovasculature. In comparison to the large body of literature of fMRI networks and brain connectivity, the evaluation of fPET resting-state networks is based on FDG-PET as a direct and quantitative index of neural metabolic activity without cerebrovascular confounds (Magistretti et al., 1999). Multi-modality imaging including simultaneous MR-PET has motivated important developments in PET detector technology, which have led to improved sensitivity for low signal detection methods including fPET (Chen et al., 2018). Future PET detector technologies may have the potential to further improve the detection sensitivity, which have the potential for the temporal resolution of functional PET to approach that of functional MRI.

### Future directions

A limitation of the FDG-fPET methodology is that the biosafety constraints on the dose of the administered radioactivity produces noisy PET images with low SNR. The differing SNR profiles between the sPET, fPET and fMRI modalities motivated the use of a percentile ‘thresholding’ approach, rather than a statistical threshold which would likely require different thresholds for each modality due to differences in statistical power. The percentile approach allowed comparison of the most highly connected regions between the modalities. We note that the atlas-based decomposition (region-of-interest) analysis of resting-state connectivity that we applied is less sensitive to the signal-to-noise limitations of fPET, and thus higher signal-to-noise timeseries data would likely be required for individual voxel-based analyses. The potential of the FDG-fPET technique is that it theoretically provides a more direct measure of neuronal activity than fMRI and does not have the susceptibility artefacts present in BOLD-fMRI. However, this advantage may not be realised in practice as the connectivity estimates in subcortical regions, which are some of the most interconnected regions in the human brain, showed low region degree in both FDG-fPET and BOLD-fMRI. This limitation may be addressed through post-processing techniques that can improve PET resolution by using synergistic MR-PET reconstruction techniques (Sudarshan et al., 2020). Future work should aim to further characterise and improve the noise properties of the fPET signal, as well as optimise the acquisition and data preparation procedures.

We encountered a technical limitation during PET post-processing that future work should address. The reconstruction software was unable to provide non-integer second fPET bins, which prevented reconstruction of the PET listmode data into images that had a time duration that was a multiple of the fMRI TR. One consequence is that a phase shift may result in the temporal synchrony of the fMRI and fPET timeseries data. Future work should consider this potential limitation at the experiment-design phase. Reconstruction of fPET frames into a multiple of the fMRI TR would also facilitate examination of temporal similarity between the two signals. An additional technical limitation is that the processing pipeline for FDG-fPET data is immature in comparison to the pre-processing procedures for BOLD-fMRI data. As a nascent technology, FDG-fPET has not had the benefit of many years of work validating acquisition parameters, data preparation and signal detection optimisation, including characterising of potential physiological artifacts that may be possible to remove with filtering techniques. Work in these fields are presently improving the radiotracer administration (e.g., (Jamadar et al., 2019a); (Rischka et al., 2018), attenuation correction (Baran et al., 2018), motion correction (Chen et al., 2019) and data analysis (Li et al., 2019) techniques.

An important distinction to make is that the brain connectivity reported here is not comparable to ‘dynamic’ connectivity as reported in the literature over the past few years. While the fPET metabolic connectivity measures are dynamic in the sense that they cross correlate the regional *time-courses* of FDG uptake over the scan period, the approach is nevertheless temporally-invariant (stationary) as the correlation is expected to be robust to temporal reordering of the measurement timepoints. In other words, the Pearson r-values describing the metabolic connectivity would be unchanged by permuting the order of the PET images. Whilst other authors have used the term ‘static’ to refer to stationary measures of functional connectivity, we do not use the term ‘static metabolic connectivity’ to avoid confusion with our use of static PET to denote across-subject PET covariance (see (Liégeois et al., 2017) for discussion of static versus dynamic functional connectivity and the mathematical basis of stationarity in haemodynamic connectivity). Our approach to analyse stationary fPET metabolic connectivity was to compare metabolic connectivity with the most common method for estimation of functional connectivity in the BOLD-fMRI literature (Liégeois et al., 2017); (Preti et al., 2017), which is the basis of our knowledge of integrative brain connectivity. However, truly ‘dynamic’ brain connectivity does take into account the temporal fluctuations of functional and metabolic connectivity, since the temporal ordering of the timeseries of brain images is important (Calhoun et al., 2014). Dynamic connectivity approaches include sliding-time window approaches and models of switching between microstates (Preti et al., 2017). Whilst it is important that future studies examine the dynamics of FDG-fPET metabolic connectivity, the current temporal resolution of fPET methodology will present a major challenge.

## Conclusion

Simultaneous neuroimaging studies that independently measure brain function are by nature challenging. Data from each imaging modality is optimally acquired using a distinctive experimental protocol and instrumentation that may adversely influence the integrity of the complementary measures of brain function. Simultaneous FDG-PET/BOLD-fMRI represents a significant advance over previous approaches. Our experimental approach to simultaneously image two mechanisms that underpin the dynamics of energy consumption in the brain via glucose metabolism and the cerebrovascular haemodynamic response, provides a unique opportunity to investigate the fundamental basis of human brain connectivity. In this study, we have reported the first FDG-fPET metabolic connectivity in humans, discovered that metabolic connectivity is more predictive of BOLD-fMRI haemodynamic connectivity than conventional FDG-PET metabolic covariance measures. Notably, these findings motivate and provide a basis for future metabolic brain connectivity studies across the human life span and investigations of novel biomarkers for neurological and neurodegenerative diseases.

## Acknowledgements

This work was supported by an Australian Research Council (ARC) Linkage Project (LP170100494) that includes financial support from Siemens Healthineers. SDJ, PGDW & GFE are supported by the ARC Centre of Excellence for Integrative Brain Function (CE140100007). SDJ is supported by an ARC Discovery Early Career Researcher Award (DE150100406) and National Health and Medical Research Council Fellowship (APP1174164).

## Author Contributions

SDJ, PGDW & GFE designed the research question, PGDW, EXL & ERO conducted the analysis, SDJ wrote the first draft of the paper with GFE and PGDW edits. All authors contributed to manuscript preparation and review. GFE, SDJ, & ZC sourced funding for the work. The authors acknowledge Richard McIntyre, Alexandra Carey, Disha Sasan and Irene Graafsma from Monash Biomedical Imaging for their assistance in acquiring data.

The authors have no competing or conflicting interests.

## Data availability

Data is available at OpenNeuro with the accession number ds002898; https://openneuro.org/datasets/ds002898/versions/1.1.0

## Supplementary Figures

**Figure S1:**
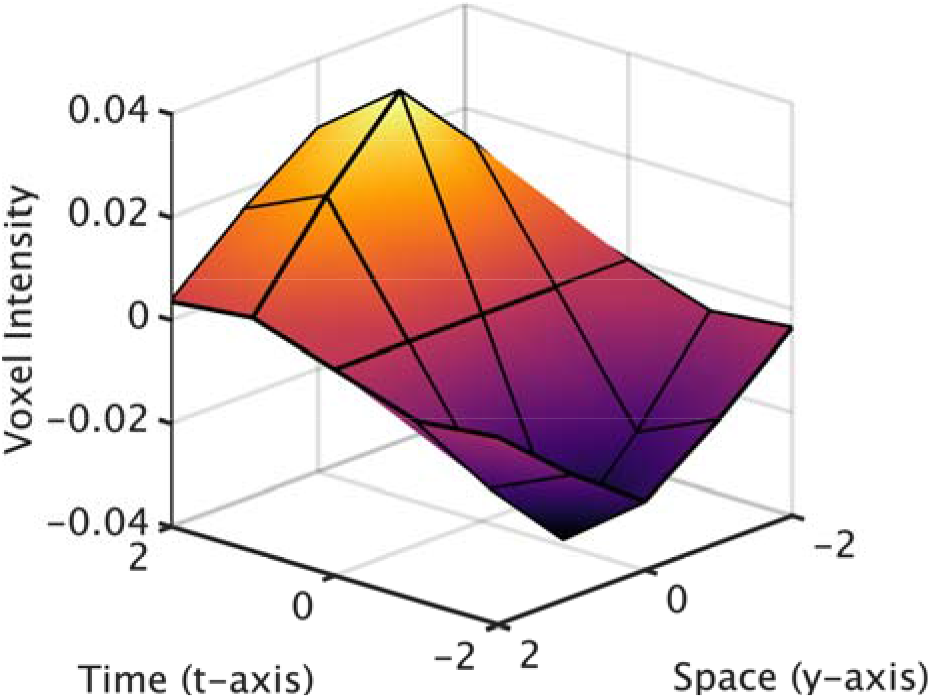
Voxel weights for the spatiotemporal filter, depicted along the y-axis and the time axis. The filter is symmetrical in the x and z axis directions.

The spatiotemporal filter is defined as the convolution of a 3-dimensional gaussian filter in the spatial domain and a 1-dimensional gaussian filter in the time domain. For the filter used in the main results of the paper, the standard deviation of the spatial gaussian was 1 voxel and of the temporal gaussian was 2 frames. The result of the spatiotemporal convolution was then modified to give negative weights on prior frames (negative time), and zero weights for the current frame.

**Figure S2:**
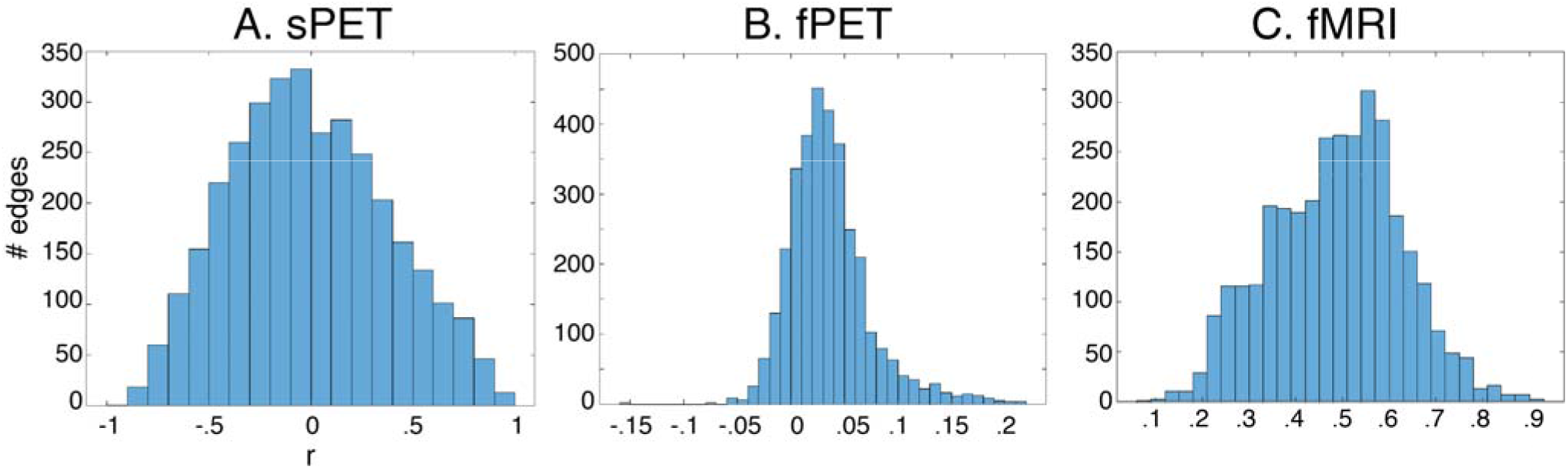
Distribution of edges across r values for each of the 3 modalities.

**Figure S3:**
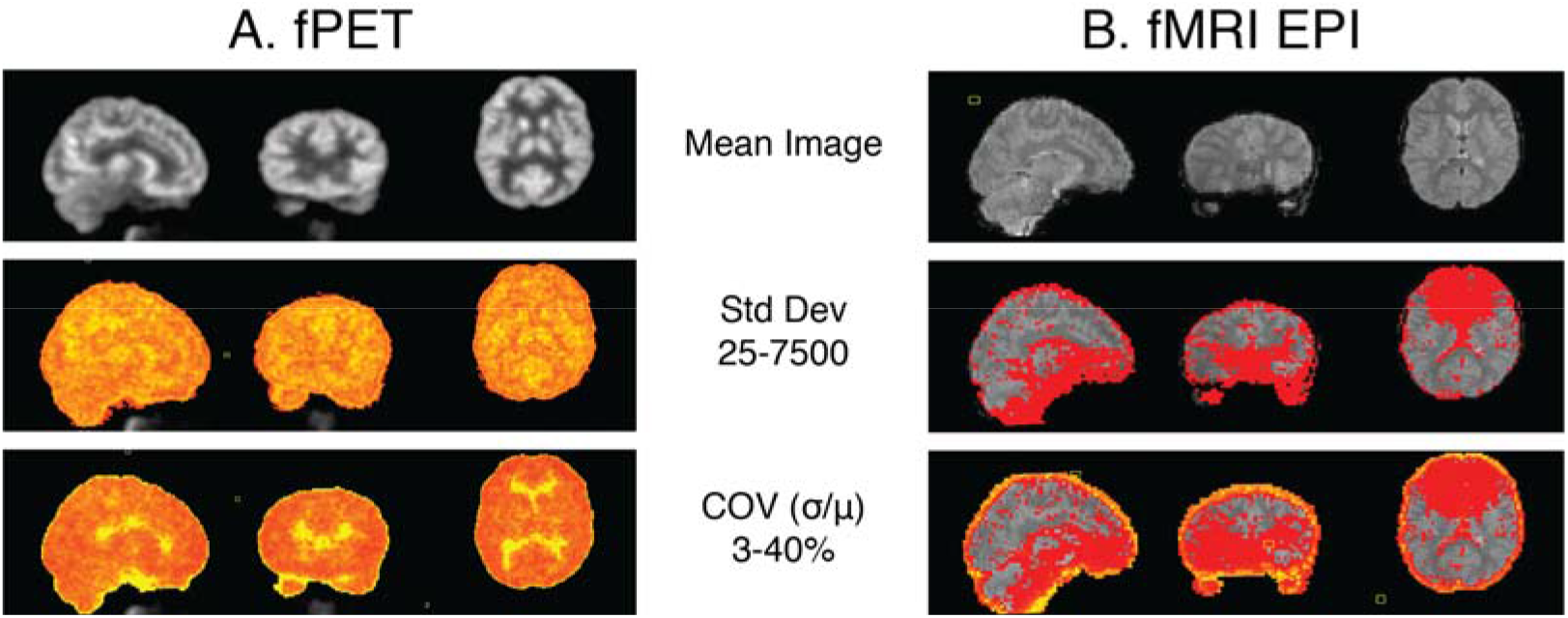
Raw images for one individual subject showing signal intensity variation across the brain. Top panel shows mean image across a 10-minute run; middle panel shows standard deviation (Std Dev) across the same run; and lower panel shows the coefficient of variation (COV, standard deviation divided by the mean) across the run.

**Figure S4.**
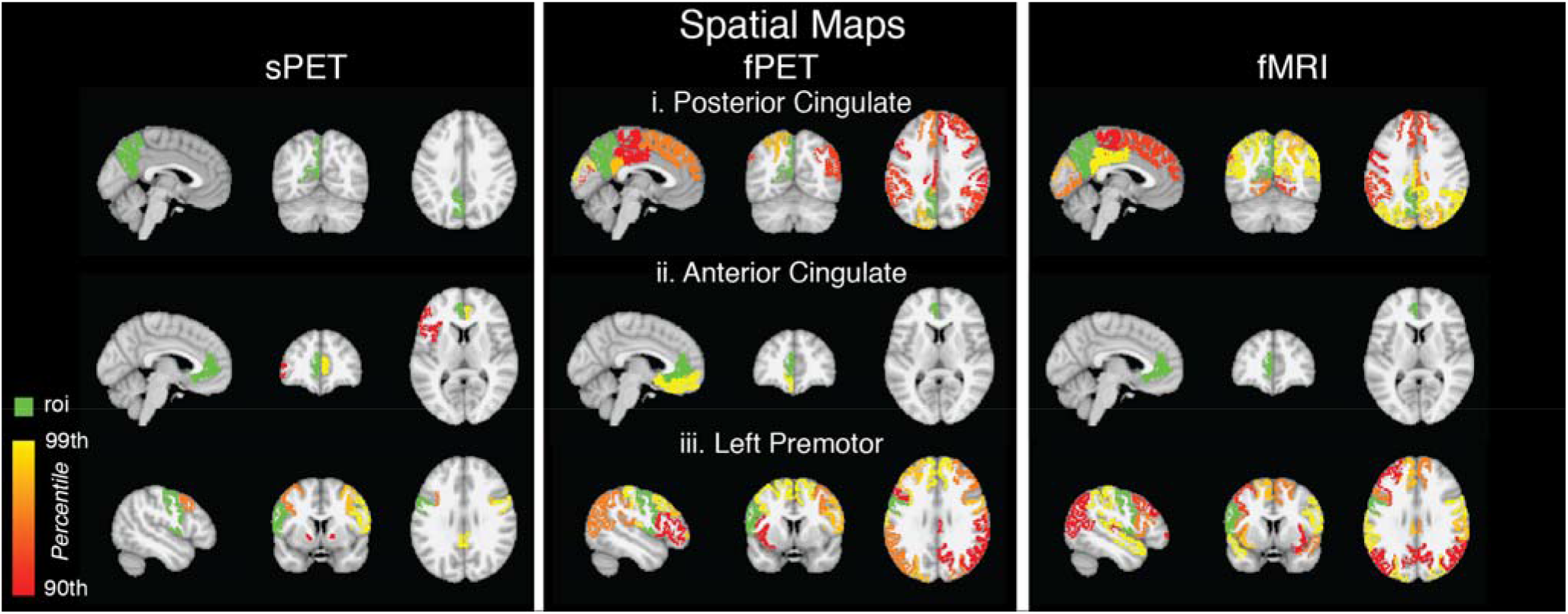
Spatial maps from three exemplar ROIs, i. the posterior cingulate, ii. anterior cingulate, and iii. left premotor cortex. These are spatial maps of the same data presented in the connectivity matrices above, and show the top 10% of regions correlated with the seed region (green).

**Figure S5.**
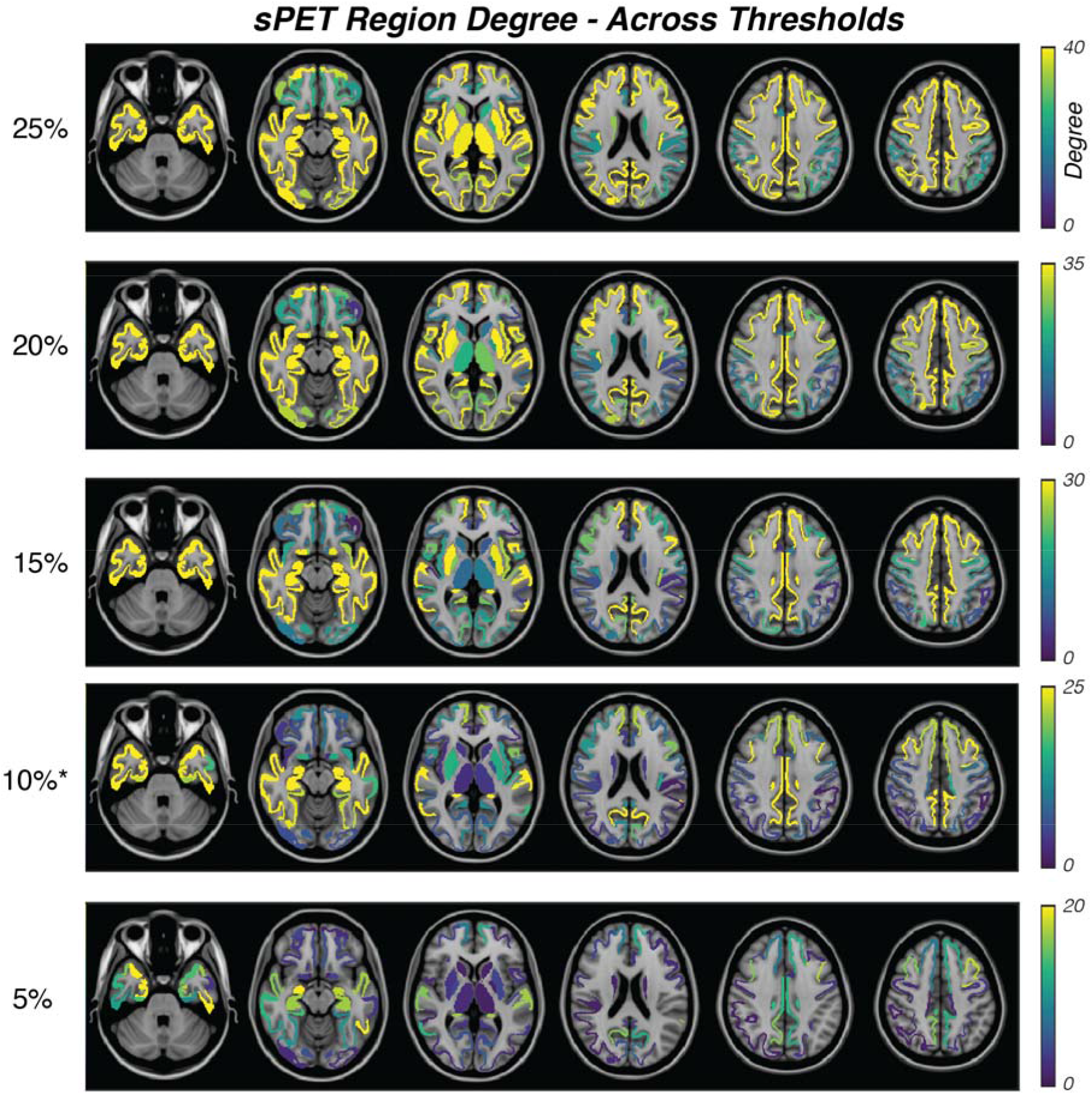
Variability of sPET region degree with different thresholds. In the manuscript, we reported the spatial distribution of the top 10% most connected regions; here, we show the spatial distribution of the top 25, 20, 15, 10, 5% most-connected nodes to illustrate how the results vary with different thresholds.

**Figure S6.**
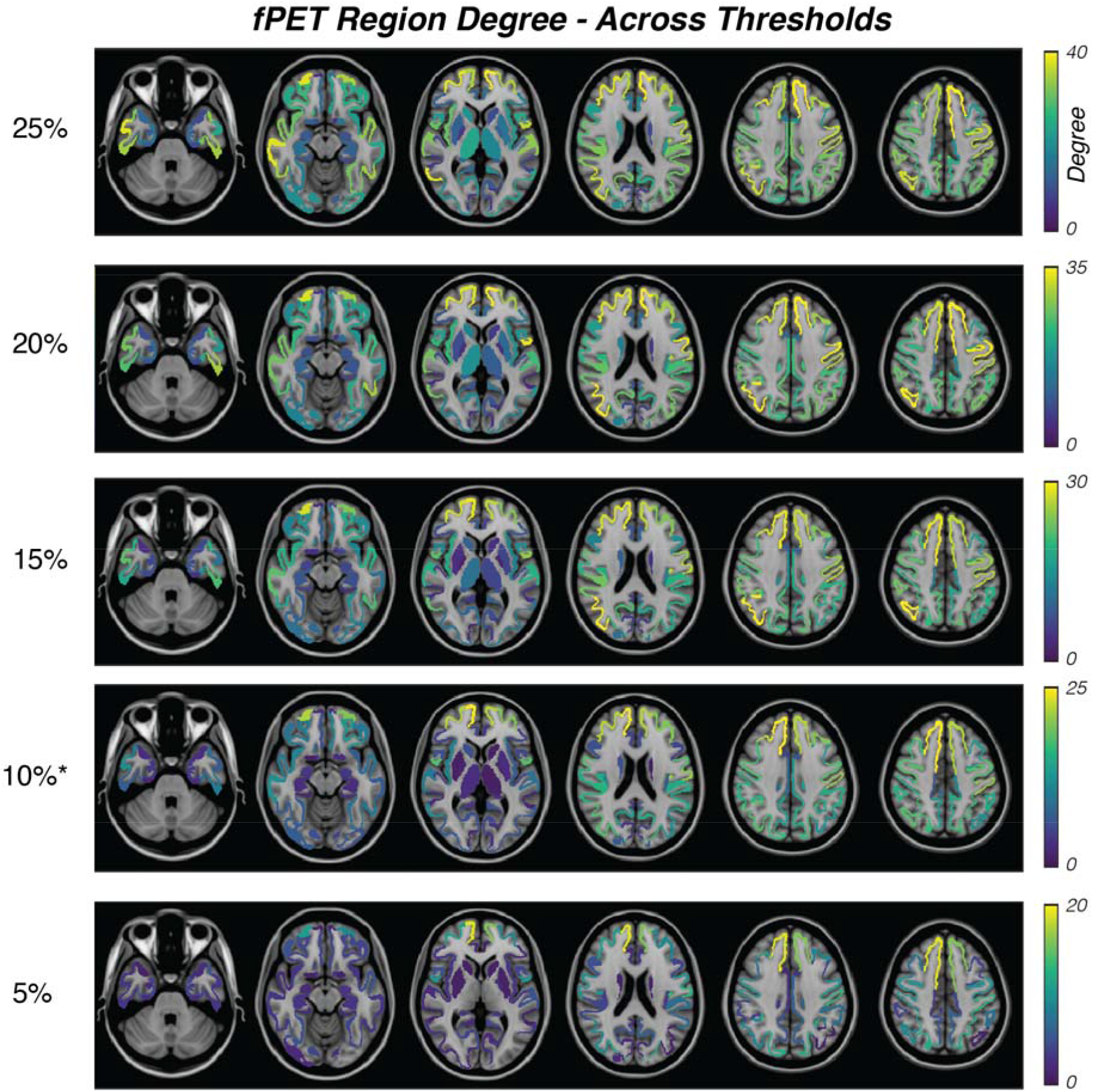
Variability of fPET region degree with different thresholds. In the manuscript, we reported the spatial distribution of the top 10% most connected regions; here, we show the spatial distribution of the top 25, 20, 15, 10, 5% most-connected nodes to illustrate how the results vary with different thresholds.

**Figure S7.**
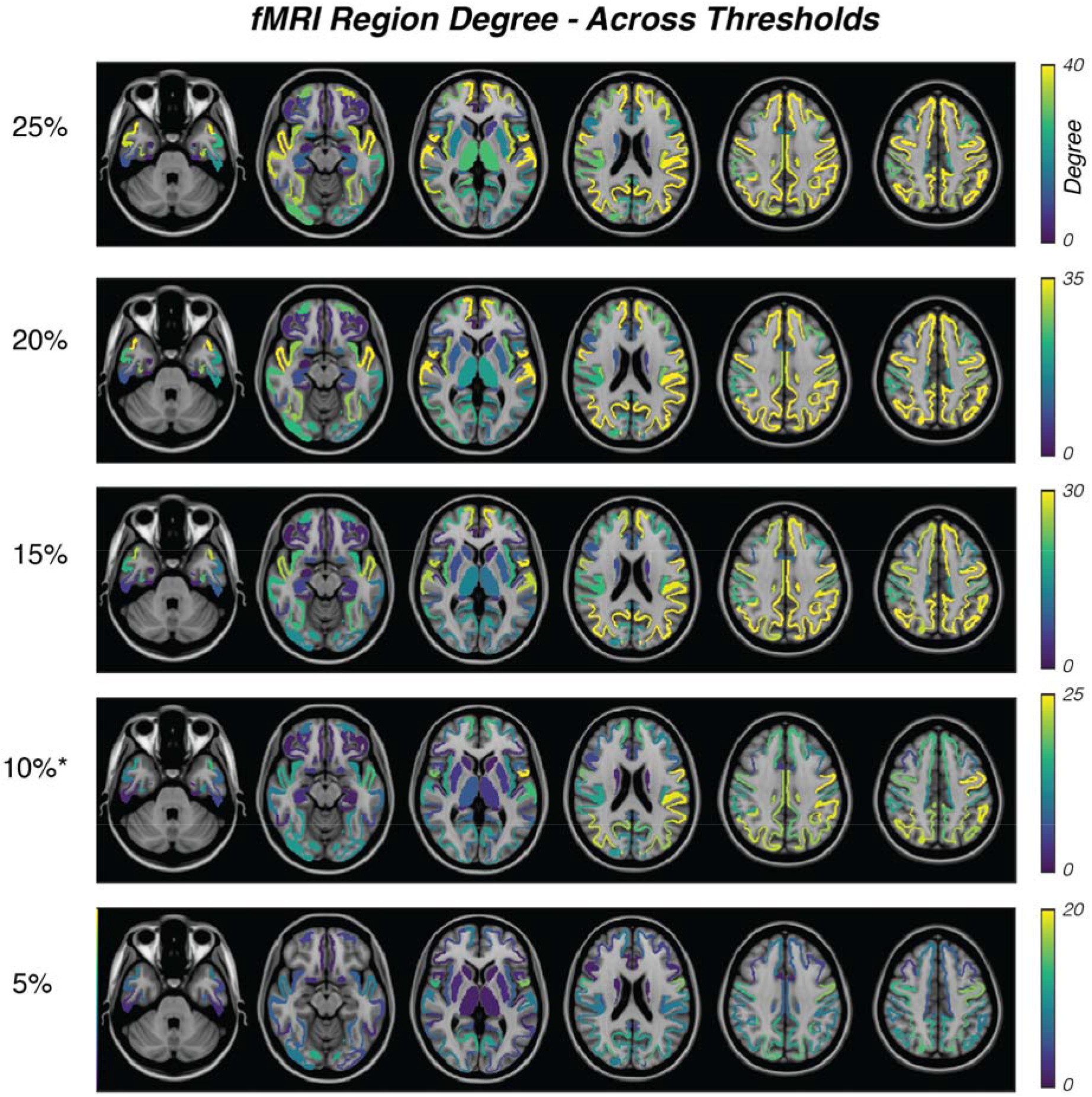
Variability of fMRI region degree with different thresholds. In the manuscript, we reported the spatial distribution of the top 10% most connected regions; here, we show the spatial distribution of the top 25, 20, 15, 10, 5% most-connected nodes to illustrate how the results vary with different thresholds.

## Notes

### Competing Interest Statement

The authors have declared no competing interest.

